# TREM2-H157Y Increases Soluble TREM2 Production and Reduces Amyloid Pathology

**DOI:** 10.1101/2021.10.04.463020

**Authors:** Wenhui Qiao, Yixing Chen, Yuka A. Martens, Chia-Chen Liu, Joshua Knight, Fuyao Li, Kai Chen, Kurti Aishe, Francis Shue, Maxwell V. Dacquel, John Fryer, Na Zhao, Guojun Bu

## Abstract

The p.H157Y variant of *TREM2* (Triggering Receptor Expressed on Myeloid Cells 2) has been reported to increase Alzheimer’s disease (AD) risk. This mutation in the extracellular domain of TREM2 localizes at the cleavage site, leading to enhanced shedding. Here, we generated a novel *Trem2* H157Y knock-in mouse model to investigate how this H157Y mutation impacts TREM2 proteolytic processing, synaptic function, and AD-related amyloid pathology. Consistent with previous *in vitro* findings, TREM2-H157Y increases the amount of soluble TREM2 (sTREM2) in the cortex and serum of mutant mice compared to the wild type controls. Interestingly, the *Trem2* H157Y variant enhances synaptic plasticity without affecting microglial density and morphology. In the presence of amyloid pathology, TREM2-H157Y surprisingly accelerates Aβ clearance and reduces amyloid burden and microgliosis. Taken together, our findings support a beneficial effect of the *Trem2* H157Y mutation in synaptic function and in mitigating amyloid pathology. Considering the genetic association of *TREM2* p.H157Y with AD, we speculate TREM2-H157Y might increase AD risk through an amyloid-independent pathway, as such its effects on tauopathy and neurodegeneration merit further investigation.

## Introduction

Alzheimer’s disease (AD) is a chronic neurodegenerative disease characterized by the pathological deposition of extracellular amyloid plaques and intraneuronal hyperphosphorylated tau tangles, as well as a prominent microglia activation responding to neuropathology and neurodegeneration (DeTure & Dickson, 2019, Guo, Zhang et al., 2020, Querfurth & LaFerla, 2010). Rare variants of multiple microglia genes are found to be associated with AD risk (Simsvan der Lee et al., 2017), including Triggering Receptor Expressed on Myeloid Cells 2 (*TREM2*). In particular, the *TREM2* p.H157Y variant was identified from a relatively small number of carriers and conferred an increased AD risk with an odds ratio (OR) of 11.01 (MAF, 0.4%) in a Han Chinese cohort (Jiang, Tan et al., 2016), whereas in a Caucasian cohort used in the Alzheimer’s Disease Sequencing Project, the OR was 4.7 (MAF, 0.06%) (Song, Hooli et al., 2017). However, how this rare *TREM2* variant impacts its function as it relates to AD risk is not clear.

TREM2 is an immunoreceptor exclusively expressed in microglia in the central nervous system and in myeloid cells (e.g., macrophage) in the periphery (Ulland & Colonna, 2018). Structurally, it consists of an Ig-like V type domain, stalk region, a transmembrane domain, and a short cytoplasmic tail (Kober, Alexander-Brett et al., 2016). Most AD-risk variants (e.g., p.R47H, p.R62H) of *TREM2* (Benitez, Cruchaga et al., 2013, Guerreiro, Wojtas et al., 2013, Jonsson & Stefansson, 2013) are located in exon2 which encodes an Ig-like domain. These pathogenic mutations often lead to ineffective binding of ligands such as Aβ oligomers (Vilalta, Zhou et al., 2021, Zhao, Wu et al., 2018, Zhong, Wang et al., 2018), fibrillar Aβ-associated anionic lipids (Wang, Cella et al., 2015), LDL (Song et al., 2017, Yeh, Wang et al., 2016), HDL (Song et al., 2017), and apolipoproteins (Atagi, Liu et al., 2015, Yeh et al., 2016). These impairments are further associated with microglial dysfunction in phagocytosis *in vitro* (Kleinberger, Yamanishi et al., 2014, Yeh et al., 2016, Yin, Liu et al., 2016) and amyloid plaques engulfment *in vivo* (Song, Joshita et al., 2018, Yuan, Condello et al., 2016). In contrast, the p.H157Y variant is located in exon3, encoding the stalk region. Intriguingly, the H157-S158 site was identified as the ADAM10/17 cleavage site that produces soluble TREM2 (sTREM2) where the H157Y mutant enhances this shedding (Feuerbach, Schindler et al., 2017, Schlepckow, Kleinberger et al., 2017, Thornton, Sevalle et al., 2017). Ectopic TREM2-H157Y expression in the HEK293 cells increases sTREM2 in conditioned medium accompanied by reduced membrane-associated mature full-length TREM2 (Schlepckow et al., 2017, Thornton et al., 2017). The increased TREM2 shedding might be related to impaired phagocytosis of pHrodo-E.Coli in HEK293 cells (Schlepckow et al., 2017) and decreased TREM2 signaling activation in response to phosphatidylserine in 2B4 T cells (Song et al., 2017). Despite these *in vitro* observations, the AD-related outcomes of *TREM2* H157Y mutation *in vivo* remain unknown.

Towards this, we generated a novel *Trem2* H157Y knock-in mouse model through CRISPR-cas9 technology. We found that TREM2-H157Y increased sTREM2 production. Moreover, TREM2-H157Y enhanced synaptic plasticity but did not affect microglial number and morphology. In the presence of amyloid pathology, TREM2-H157Y reduced amyloid burden, toxic Aβ oligomer, and microgliosis. Our results imply that the TREM2-H157Y might be beneficial to brain function and in reducing amyloid pathology and related toxicity.

## Results

### Generation of *Trem2* H157Y knock-in mouse model

TREM2-H157 is located where TREM2 undergoes shedding to produce sTREM2 (Fig 1A) (Feuerbach et al., 2017, Schlepckow et al., 2017, Thornton et al., 2017). To study the *in vivo* effects of the *Trem2* H157Y mutation, we introduced a C>T substitution in exon3 through CRISPR/Cas9 technology to create the missense H157Y mutation (Fig 1B). Two founders (1^#^ and 2^#^) were obtained with no off-target mutation observed in the offspring of either founder (Fig EV1A and B). Results reported below were generated using the offspring of Founder 1^#^ unless otherwise stated. By crossing the *Trem2* H157Y heterozygous mice, we obtained three genotypes: wild type (*Trem2*^*+/+*^, referred to as WT), heterozygous (*Trem2*^*+/H157Y*^, referred to as Het), and homozygous (*Trem2*^*H157Y/H157Y*^, referred to as Hom). Littermates of the three genotypes were used to investigate the impact of the *Trem2* H157Y mutation.

**Figure 1.**
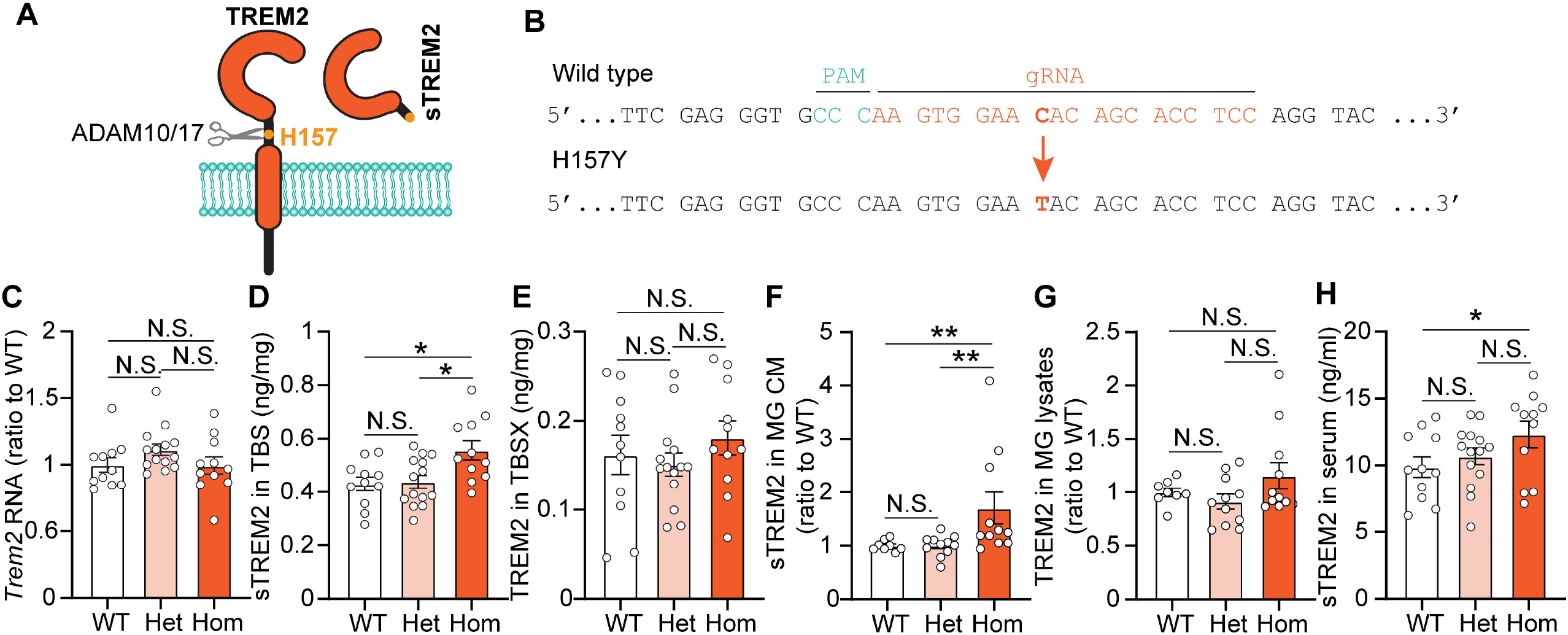
*Trem2* H157Y mutation increases sTrem2. A Schematic illustration of sTrem2 generation. B Trem2 H157Y knock-in mice were generated by introducing a C>T mutation (bold orange) via CRIPR-Cas9. Protospacer region recognized by guide RNA (gRNA) is shown in orange. Protospacer adjacent region (PAM) is shown in green. C Trem2 mRNA level was examined in the cortex of mice at 6 months of age. N =11-14 mice per genotype, mixed sex. D-E TREM2 level was examined by ELISA in cortical extracts obtained by sequential protein extraction with TBS (C) and TBSX(D) from mice at 6 months of age. N =11-14 mice per genotype, mixed sex. F-G TREM2 level was examined by ELISA in conditioned medium (CM) (E) and RIPA lysates (F) of primary microglia (MG). TREM2 amount was normalized to the total protein level of cell lysates followed by another normalization to the values of WT littermates. N=8-11 pups per genotype. H TREM2 level in the serum of mice at 6 months of age was examined by ELISA. N =11- 14 mice per genotype, mixed sex. Data information: Data are presented as Mean±SEM. Kruskal-Wallis tests with uncorrected Dun’s multiple comparisons were used in C-H. N.S., not significant. * p<0.05. **p<0.01.

### TREM2-H157Y increases the production of sTREM2

At the transcription level, there was no significant change of cortical *Trem2* mRNA level in *Trem2* H157Y Het or Hom mice compared to WT mice at 6 months of age (Fig. 1C) compared to the WT mice. To evaluate TREM2 protein levels, proteins were sequentially extracted from cortex with Tris-buffered saline (TBS) and TBSX (TBS+1% Triton X-100) and analyzed by N-terminal TREM2-capturing ELISA. Although membrane bound TREM2 in TBSX did not differ between genotypes (Fig. 1E), there was an increase of sTREM2 in the TBS lysates in Hom compared to Het and WT mice (Fig. 1D).

To further examine TREM2 processing in microglia, we cultured cortical primary microglia from Het breeder littermate pups. Consistent with in vivo findings, we observed an increase of sTREM2 in conditioned medium (CM) from Hom microglia compared to Het and WT microglia (Fig 1F). The membrane associated TREM2 in microglia RIPA lysates did not differ between genotypes (Fig 1G). Further supporting an increase of sTREM2 production by the *Trem2* H157Y mutation, we observed higher levels of serum sTREM2 in Hom mice compared to WT and Het mice (Fig. 1H). Together, our results support an effect of the *Trem2* H157Y mutation on increasing sTREM2 production in homozygous mice which are consistent with prior *in vitro* findings (Schlepckow et al., 2017, Thornton et al., 2017).

### TREM2-H157Y does not affect microglia density and morphology

To quantify the microglia density and assess the morphology of microglia, we performed IBA1 immunofluorescence staining of brain slices from *Trem2* H157Y knock-in mice at 6 months of age. Microglia density and cell body size did not change with the *Trem2* H157Y mutation (Fig EV2A-C). Analyses after microglia skeletonization (EV2D-F) showed no significant differences in the branch number, junction number, or total branch length per microglia between genotypes (EV2G-I). These results suggest TREM2**-**H157Y does not affect microglia density and morphology *in vivo* under physiological conditions.

### TREM2-H157Y enhances synaptic plasticity

It has been reported that microglia play important roles in synaptic pruning and neural circuit regulation (Filipello, Morini et al., 2018). Thus, we assessed whether the *Trem2* H157Y mutation affects synaptic plasticity. We performed hippocampal long-term potentiation (LTP) in WT and Hom mice at 6 months of age. While the basic transmission and presynaptic facilitation were unaffected (Fig 2A and B), we observed an enhanced LTP in the Hom mice compared to WT mice (Fig 2C and D).

**Figure 2.**
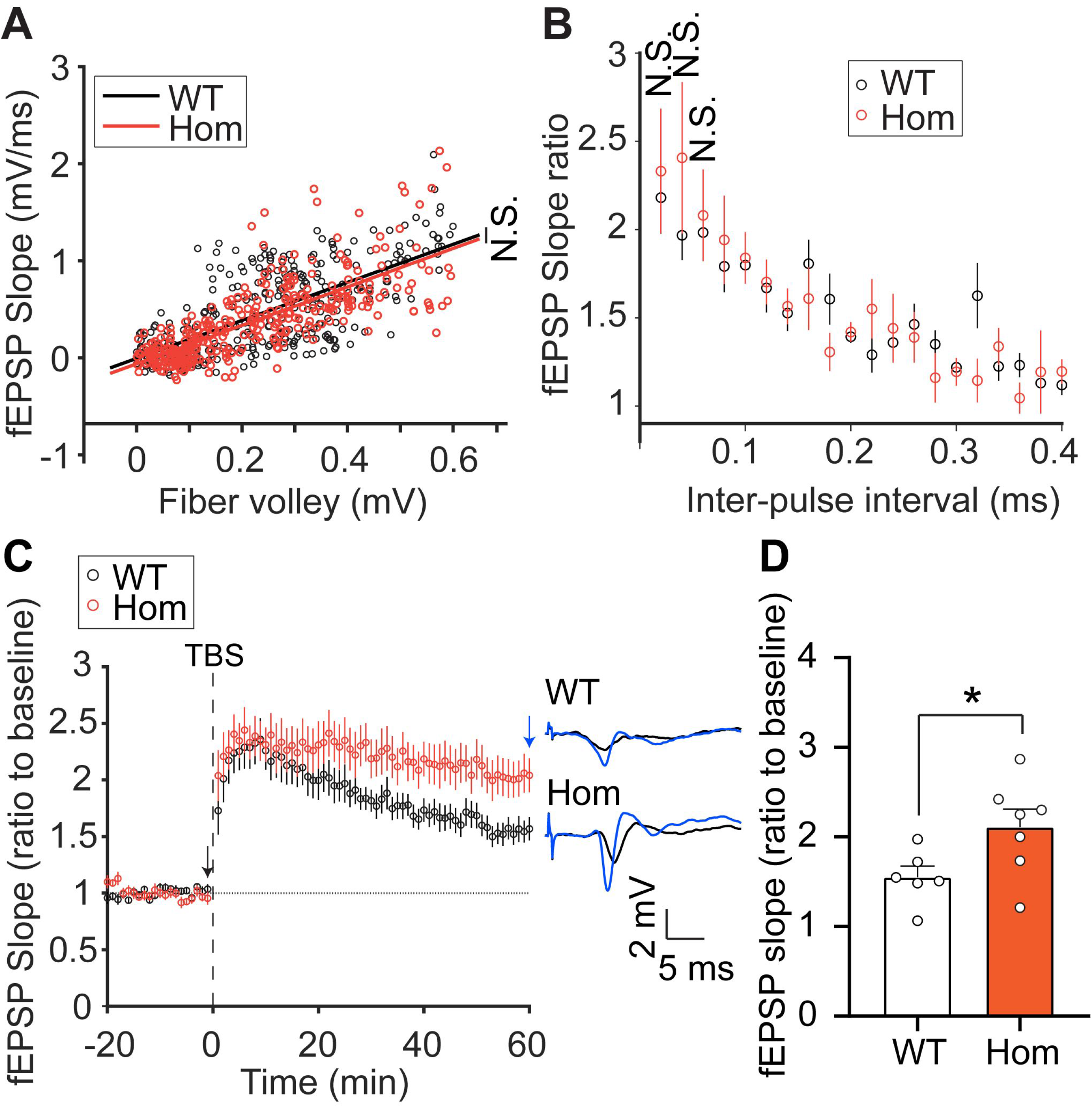
*Trem2* H157Y mutation enhances synaptic plasticity. A The input-output curves for WT and Hom mice at 6 months of age are shown as linear regressions of fEPSP slopes in response to pre-synaptic fiber volley amplitudes. n = 13 brain slices from 6-7 male mice/group. B Paired-pulse facilitation (PPF) profiles were achieved with increased inter-pulse intervals (IPI) are shown. n = 11-14 brain slices from 6-7 male mice/group. C-D Theta-Burst Stimulus (TBS) induced LTP profiles for WT and Hom mice are shown as averaged and normalized fEPSP slopes (C). Example recording traces before and after TBS stimulation are shown. The averages fEPSP slope in the last five minutes are compared between groups (D). n = 12 brain slices from 6-7 male mice/group. Data information: Data are presented as Mean±SEM. One-Way ANOCOVA with comparison of slopes was used in A. Wilcoxon Rank-sum tests were used in B. Unpaired t test was used in D. N.S., not significant. *p < 0.05.

To examine whether this strengthened synaptic capability is correlated with enhanced cognitive performance, we conducted a battery of behavioral tests with *Trem2* H157Y knock-in mice. We did not observe significant performance differences in anxiety (Fig EV3A) and associative memory assessments (Fig EV3C and D) between genotypes. However, using Y-maze spontaneous tests, we observed a trending performance improvement of spatial working memory in Hom mice compared to Het mice while no difference between Het mice and WT mice (Fig EV3B; Het vs Hom, p = 0.06). These results together support a beneficial effect of TREM2-H157Y on synaptic plasticity, even though it did not translate into significant enhancement at the behavioral level.

### TREM2-H157Y reduces amyloid burden in 5xFAD mice

To investigate the effects of H157Y mutation on AD-related amyloid pathogenesis, *Trem2* H157Y knock-in mice were crossed with 5xFAD amyloid model mice to generate littermates with three genotypes, 5xFAD/WT (*5xFAD; Trem2*^*+/+*^), 5xFAD/Het (*5xFAD; Trem2*^*+/H157Y*^), and 5xFAD/Hom (*5xFAD; Trem2*^*H157Y/H157Y*^). Animals were harvested at 8.5 months of age to assess amyloid pathology at a middle-to-late stage of amyloid development in the cortex (Jay, Hirsch et al., 2017).

Total Aβ immunostaining with MOAB2 antibody revealed significant reductions plaque number (Fig 3A and B) in 5xFAD/Hom mice compared to 5xFAD/WT mice. Plaques from all three genotypes were found to be similar in size (Fig 3C). We did not observe significant decreases of the X34-positive fibrillar Aβ signal with the *Trem2* H157Y mutation (Fig EV4A-C). Moreover, we detected Aβ40 and Aβ42 by ELISA in cortical lysates obtained through sequential TBS, TBSX, and guanidine (GND) extraction. Consistent with the reduction of total Aβ in staining, we observed significant reductions of Aβ40 and Aβ42 in GND lysates (Fig 3D and E) from 5xFAD/Hom mice compared with 5xFAD/WT mice. The 5xFAD/Het group exhibited no significant differences compared to 5xFAD/Hom and 5xFAD/WT groups (Fig 3D and E). We did not observe a significant decrease of Aβ40 and Aβ42 in both TBS and TBSX lysates with the *Trem2* H157Y mutation (Fig EV4D-G).

**Figure 3.**
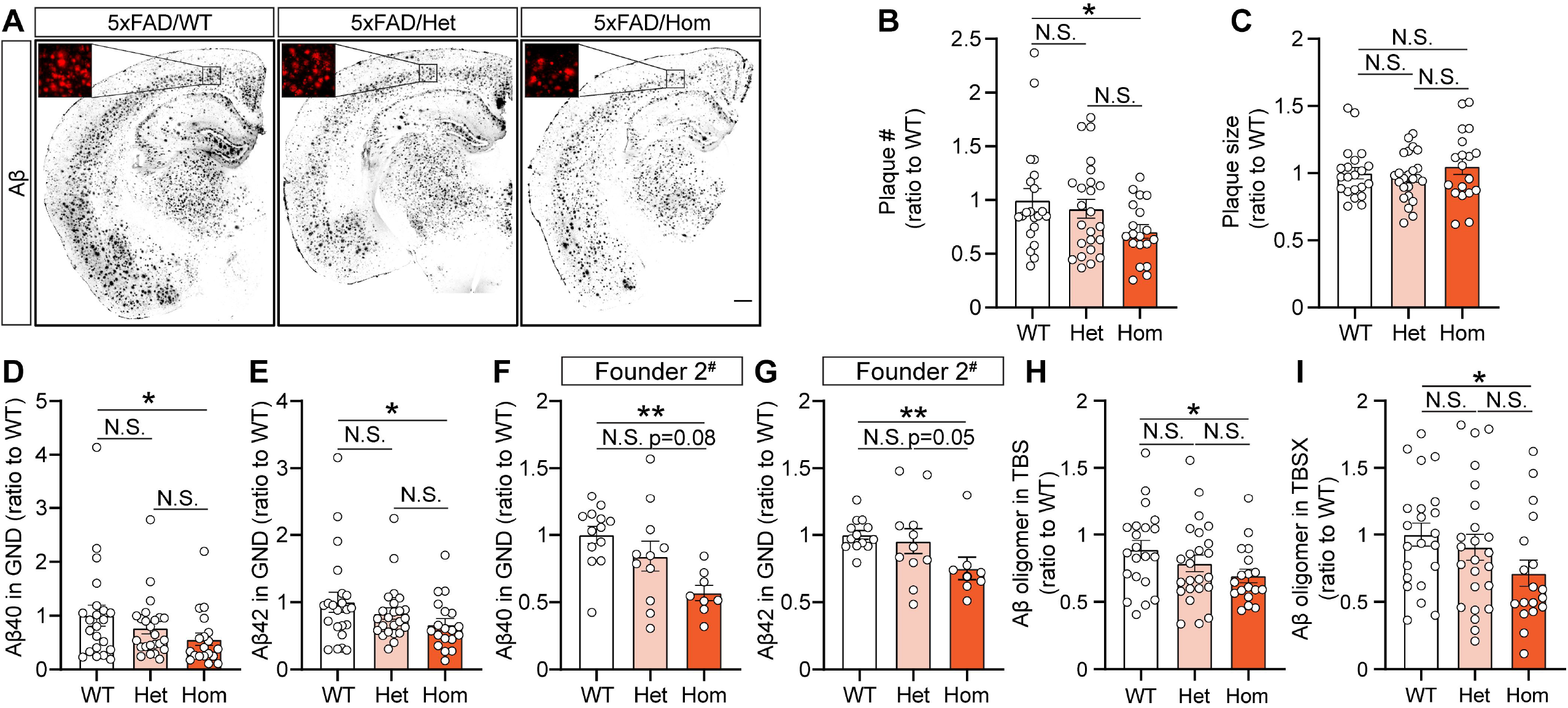
*Trem2* H157Y mutation leads to reduction in amyloid in 5xFAD mice. A Representative images of amyloid staining are shown for 5xFAD/WT, 5xFAD/Het, and 5xFAD/Hom mice at the age of 8.5 months of age. Scale, 400 µm. B-C Amyloid plaque number (B) and size (C) are quantified for each genotype from mice at the age of 8.5 months of age. N =19-24 mice per genotype, mixed sex. D-E Aβ40 (D) and Aβ42 (E) are measured by ELISA in cortical guanidine lysates (GND) for each genotype. N = 19-24 mice per genotype, mixed sex. F-G Aβ40 (F) and Aβ42 (G) are measured by ELISA in cortical guanidine lysates (GND) for each genotype from Founder 2# offspring. N = 8-13 mice per genotype, mixed sex. H-I Aβ oligomer is measured by ELISA in cortical extracts obtained sequentially by TBS (H) and TBSX (I) buffer. N =19-24 mice per genotype, mixed sex. Data information: Data are presented as Mean±SEM. Kruskal-Wallis tests with uncorrected Dun’s multiple comparisons were used in B- I. N.S., not significant. * p<0.05. **p<0.01.

To confirm TREM2-H157Y effects on amyloid burden, we crossed 5xFAD mice with our second founder of the *Trem2* H157Y knock-in mice (Founder 2^#^). Consistent with the results from the Founder 1^#^ offspring, we observed significant reductions of Aβ40 and Aβ42 levels in GND lysates from 5xFAD/Hom mice compared to 5xFAD/WT (Fig 3F and G). Also, 5xFAD/Hom group showed trending decreases compared 5xFAD/Het group (Fig 3F and G; Aβ40, Het vs Hom, p=0.08; Aβ42, Het vs Hom, p=0.05). In this cohort, both TBS-Aβ40 and TBS-Aβ42 in 5xFAD/Hom mice reduced significantly compared to 5xFAD/WT and trended toward reductions compared to 5xFAD/Het mice (Fig EV4H and J; Aβ40, Het vs Hom, p=0.05; Aβ42,Het vs Hom, p=0.05). TBSX-Aβ40 was significantly reduced in 5xFAD/Het mice and trended toward a reduction in 5xFAD/Hom mice compared to 5xFAD/WT mice (Fig EV4I; WT vs Hom, p=0.07). No significant reductions of TBSX-Aβ42 were observed with the *Trem2* H157Y mutation (Fig EV4K).

We further measured the levels of Aβ oligomers, the neuronal toxic species (Walsh, Klyubin et al., 2002, Wei, Nguyen et al., 2010) in TBS and TBSX fractions, and found significant reductions in 5xFAD/Hom mice compared to 5xFAD/WT mice (Fig 3H and I). There were no significant differences between the 5xFAD/Het group and the other two groups (Fig 3H and I). We then examined Aβ toxicity-related dystrophic neurites through lysosome-associated membrane protein (LAMP1) immunostaining. We did not observe significant changes in LAMP1 signal with the *Trem2* H157Y mutation (Fig EV4L and M). Taken together, the *Trem2* H157Y mutation reduced insoluble Aβ levels and total amyloid burden in homozygous mice.

### TREM2-H157Y facilitates Aβ clearance in 5xFAD mice

To address the potential mechanism of amyloid reduction in the *Trem2* H157Y mice, we examined the APP processing products (Chen, Xu et al., 2017) and found no significant changes in the levels of sAPPα, sAPPβ, and CTFβ between groups (Fig EV5A-C), suggesting unaltered Aβ production. We then conducted *in vivo* microdialysis with awake, free-moving mice at 3 months of age (Cirrito, May et al., 2003, Liu, Zhao et al., 2017) to analyze Aβ42 clearance in the interstitial fluid (ISF) while Aβ production was inhibited with *γ*-secretase inhibitor, LY411575 (Fig 4A). The elimination kinetic analysis showed enhanced clearance of Aβ42 with decreased Aβ42 levels four hours post drug administration (Fig 4B) and a 50% reduction of Aβ42 half-life (Fig 4C) in 5xFAD/Hom mice compared to 5xFAD/WT mice.

**Figure 4.**
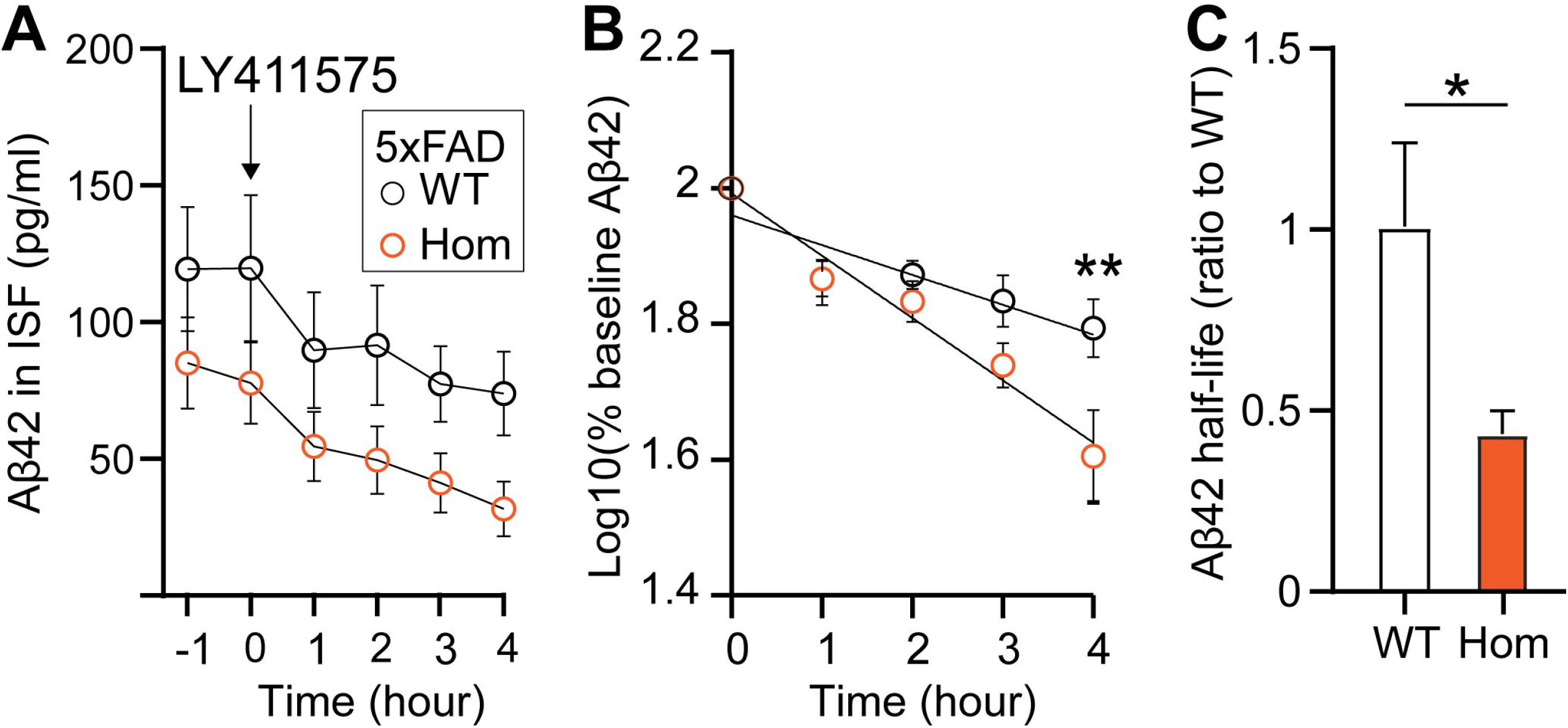
*Trem2* H157Y mutation accelerates the Aβ clearance in 5xFAD mice. A Aβ42 level is quantified by ELISA in the interstitial fluid (ISF) obtained in microdialysis experiments with WT and Hom mice at 3 months of age. N = 6-7 mice per genotype, mixed sex. At time 0, γ-secretase inhibitor LY411575 was administrated to stop the Aβ production. B Semilog plot is performed from time 0 to analyze the half-life of Aβ42 clearance. C Half-life is quantified and plotted in WT and Hom group with a normalization to WT. N = 6-7 mice per genotype, mixed sex. Data information: Data are presented as Mean±SEM. Unpaired *t* tests were used in B. Welch’s *t* test was used in C. N.S., not significant. * p<0.05. **p<0.01.

### TREM2-H157Y reduces microgliosis in 5xFAD mice

To assess the microglial responses to amyloid pathology with TREM2-H157Y, we performed immunostaining of IBA1 and the phagocytic marker CD68 in cortical brain slices from mice at 8.5 months of age. A significant reduction of microgliosis was observed with both IBA1 (Fig 5A and B) and CD68 (Fig 5D and E) signals in 5xFAD/Hom mice compared to 5xFAD/WT. The 5xFAD/Het group showed no significant differences compared to 5xFAD/WT group and 5xFAD/Hom mice. We further found positive correlation between either IBA1 or CD68 signals and Aβ42 in GND lysates, suggesting that the decreased microgliosis is likely due to a reduction in amyloid load in mice with the *Trem2* H157Y mutation (Fig 5C and F).

**Figure 5.**
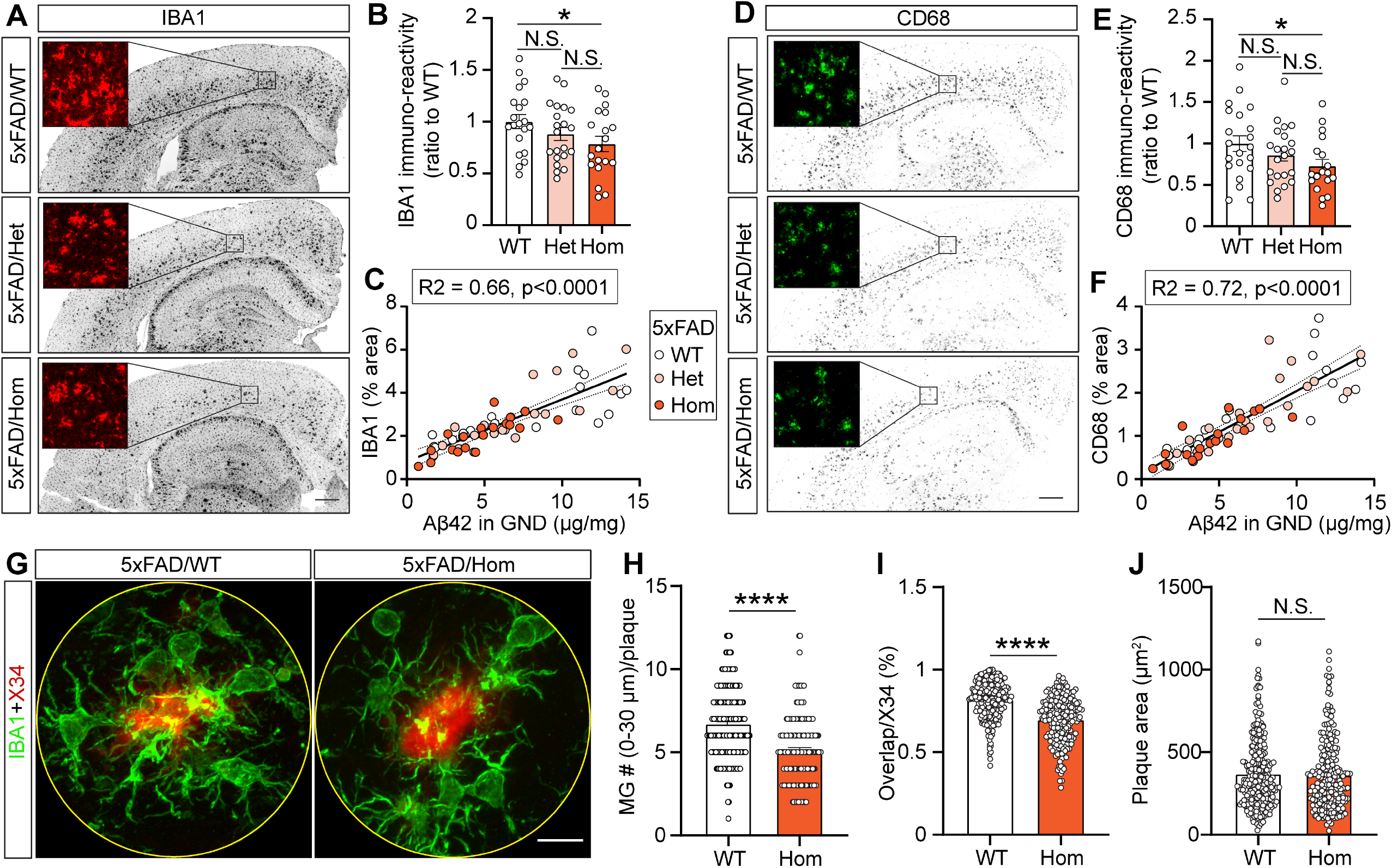
*Trem2* H157Y mutation reduces microgliosis in 5xFAD mice. A Representative images of IBA1 staining are shown for 5xFAD/WT, 5xFAD/Het, and 5xFAD/Hom mice at 8.5 months of age. Scale, 400 µm. B IBA1 immuno-reactivity is quantified for each genotype at the age of 8.5 months of age. N =19-24 mice per genotype, mixed sex. C Correlation analysis between IBA1 signals (area %) and Aβ42 in GND lysates with R2 and p value shown. N =19-24 mice per genotype, mixed sex. D Representative images of CD68 staining are shown for 5xFAD/WT, 5xFAD/Het, and 5xFAD/Hom mice at the age of 8.5 months of age. Scale, 400 µm. E CD68 immuno-reactivity is quantified for each genotype at the age of 8.5 months of age. N =19-24 mice per genotype, mixed sex. F Correlation analysis between CD68 signals (area %) and Aβ42 in GND lysates with R2 and p value shown above. N =19-24 mice per genotype, mixed sex. G Representative confocal images of IBA1 and X34 co-staining are shown for 5xFAD/WT and 5xFAD/Hom mice at the age of 8.5 months of age. Scale, 10 µm. H Microglia cell body number surrounding X34 signal is counted within a radius of 30 um for each genotype at the age of 8.5 months of age. N = 4 mice per genotype, n = 32-69 plaques/mouse. I Plaque area (X34) coverage by microglia (MG) (IBA1) is quantified for each genotype at the age of 8.5 months of age. N = 4 mice per genotype, n = 32-69 plaques/mouse. J Each plaque size is quantified for each genotype at the age of 8.5 months of age. N = 4 mice per genotype, n = 32-69 plaques/mouse. Data information: Data are presented as Mean±SEM. Kruskal-Wallis tests with uncorrected Dun’s multiple comparisons were used in B and E. Wilcoxon Rank-sum tests were used in H-J. N.S., not significant, * p<0.05. ****p<0.0001.

Plaque-associated microglia have been identified as a critical pathological event in response to amyloid (DeTure & Dickson, 2019). We found that the number of microglia associated with amyloid plaques (Fig 5G and H) and plaque area coverage by microglia (Fig 5I) were significantly reduced in 5xFAD/Hom mice compared to 5xFAD/WT mice. Plaque size did not differ between 5xFAD/Hom and 5xFAD/WT mice (Fig 5J). Overall, we observed reduced microgliosis in 5xFAD mice with the *Trem2* H157Y mutation.

## Discussion

In this study, we provided *in vivo* evidence that TREM2-H157Y promotes TREM2 shedding in our novel *Trem2* H157Y knock-in mouse models. Moreover, we found TREM2-H157Y enhances synaptic plasticity, facilitates Aβ clearance, and reduces amyloid burden.

Consistent with previous *in vitro* findings (Schlepckow et al., 2017, Thornton et al., 2017), we observed significantly higher sTREM2 level in cortical TBS lysate, conditioned medium of primary microglia, and peripheral serum from mice with the *Trem2* H157Y mutation. We did not observe significant changes in membrane associated full-length TREM2. Since N-terminal TREM2 ELISA does not distinguish mature and immature full length TREM2, we cannot conclude that TREM2-H157Y specifically reduces mature TREM2 in our mouse model as described in the *in vitro* studies (Schlepckow et al., 2017, Thornton et al., 2017).

*Trem2* p.R47H and p.Y38C variants impair synaptic plasticity through a loss of TREM2 function (Jadhav, Lin et al., 2020, Ren, Yao et al., 2020). In contrast, we observed enhanced synaptic plasticity in *Trem2* H157Y knock-in mice implying there might be a different mechanism by which TREM2-H157Y affects brain functions compared to other variants such as p.R47H or p.Y38C. Considering the increased sTrem2 in our *Trem2* H157Y mice, we speculate that the enhancement of synaptic plasticity may be due to the increased levels of sTREM2. Such a mechanism would be consistent with previous findings that exogenous sTREM2 enhances LTP in an amyloid mouse model (Zhong, Xu et al., 2019). Additionally, it has been reported that sTrem2 is associated with neurons (Song et al., 2018) further implicating sTREM2 may affect synaptic function. However, elucidating the roles of TREM2 and sTREM2 in regulating neuronal activity needs more comprehensive studies.

The mechanism by which TREM2-H157Y facilitates Aβ clearance and lowers amyloid burden is not well understood. However, we speculate that this might link to the interaction between sTREM2 and Aβ. It has been shown that the Aβ oligomer can bind to TREM2 or sTREM2 (Lessard, Malnik et al., 2018, Vilalta et al., 2021, Zhao et al., 2018, Zhong et al., 2018). Also, Aβ oligomers stimulate sTREM2 production in a dose-dependent manner *in vitro* and sTREM2 in return inhibits Aβ aggregation (Vilalta et al., 2021), suggesting that sTREM2 could facilitate Aβ diffusion and clearance *in vivo*. Studies have shown that elevating sTREM2 through exogenous administration or AAV-mediated overexpression significantly reduces amyloid burden (Zhong et al., 2019). Depleting microglia abolishes the rescuing effect of sTREM2, suggesting that sTREM2 may reduce amyloid load through microglial activation (Zhong et al., 2019). Thus, in our mouse models, increased sTREM2 by TREM2-H157Y may accelerate Aβ clearance and/or microglia activation, leading to the overall decrease of amyloid burden and related microgliosis. Microgliosis reduction may also slow down amyloid progression since phagocytic microglia with Aβ aggregates may serve as a source of seeding for amyloid plaques (Fuhrmann, Bittner et al., 2010).

Studies on *Trem2* p.R47H reveal a loss of TREM2 function in ligand binding, signaling, and microglial responses to pathological cues (Song et al., 2017, Song et al., 2018), which inspired the development of TREM2 activating antibodies to alleviate AD pathology. TREM2 antibody administration in amyloid mouse models has been found to boost microglial responses to Aβ, reduce amyloid load, toxicity, and behavioral impairments (Cheng, Danao et al., 2018, Fassler, Rappaport et al., 2021, Schlepckow, Monroe et al., 2020, Wang, Mustafa et al., 2020). While TREM2 activating antibodies stabilize the membrane form of TREM2 and related signaling, the levels of sTREM2 in serum and CSF decrease accordingly in a dose dependent manner in mice and humans (Fassler et al., 2021, Schlepckow et al., 2020, Wang et al., 2020). These findings emphasize the critical role of membrane bound TREM2 in cell-autonomous microglia activation and phagocytosis to reduce amyloid pathology. Using *Trem2* H157Y knock-in mouse models, our data alternatively suggests non-cell autonomous benefits of sTREM2 on neuronal function and Aβ clearance, encouraging a consideration of increasing sTREM2 as a potential therapeutic strategy to treat AD. Combination therapy by activating TREM2 signaling and elevating sTREM2 level should also be considered.

In summary, our study confirmed increased shedding of TREM2-H157Y *in vivo* and defined beneficial effects of TREM2-H157Y in brain function and in reducing amyloid pathology. However, these findings conflict with the genetic studies showing the increased AD risk associated with *TREM2* p.H157Y. Considering that no animal model fully mimics the AD related pathologies and 5xFAD mice merely develop amyloid pathology which recapitulates the very early stage of AD (McDade, Llibre-Guerra et al., 2021), our current data cannot address how *TREM2* p.H157Y affects late stage AD pathologies including tauopathy and neurodegeneration. Thus, more investigations are necessary to further elucidate the effect of *TREM2* H157Y mutation on AD pathogenic events, in particular the tau pathology and related neurodegeneration.

## Materials and Methods

### Generation, genotyping, and off-target analysis of *Trem2* H157Y knock-in mice

*Trem2* H157Y knock-in mice were generated via CRISPR/Cas9 by the Hope Center Transgenic Vectors Core of the Washington University (Ran, Hsu et al., 2013). CRISPR gRNAs for *in vitro* testing were identified using CRISPOR (http://crispor.tefor.net/) and synthesized as gBlocks (IDT) with the sequence 5’GGAGGTGCTGTgTTCCACTT3’. *In vitro* target specific gRNA cleavage activity was validated by transfecting N2A cells with PCR amplified gRNA gblock and Cas9 plasmid DNA (px330, addgene) using ROCHE Xtremegene HP. Cell pools were harvested 48 hours later for genomic DNA prep, followed by sanger sequencing of PCR products spanning the gRNA/Cas9 cleavage site, and TIDE analysis (https://tide.nki.nl/) of sequence trace files. CRISPR sgRNA (IDT, 20 ng/ul) and Cas9 (IDT, 50ng/ul) proteins were complexed to generate the ribonucleoprotein (RNP) for injection along with a 200 nucleotide ssODN donor DNA (synthesized by IDT, 20 ng/ul), 5’tatatcttgtcctttgctgatctgtttgccctgggacctccatccgcagtcactgccagggggtctaagaagggaccactactgtacCTGGAGGTGCTGTaTTCCACTTGGGCACCCTCGAAACTCGATGACTCCTCGGGGACCCAGAGATCTCCAGCATCTTGGTCATCTAGAGGGTctgtaatagacaaaccatgagg3’. All animal experiments were approved by institutional IACUC protocols. B6/CBA F1 mice at 3-4 weeks of age (JAX Laboratories, Bar Harbor ME, USA) were superovulated by intraperitoneal injection of 5 IU pregnant mare serum gonadotropin, followed 48 hours later by intraperitoneal injection of 5 IU human chorionic gonadotropin (PMS from SIGMA, HGC from Millipore USA). Mouse zygotes were obtained by breeding B6/CBA stud males with superovulated B6/CBA females at a 1:1 ratio. One-cell fertilized embryos were injected into the pronucleus and cytoplasm of each zygote. Microinjections and mouse transgenesis experiments were performed as described previously (Behringer, Gertsenstein et al., 2014, Pease & Saunders, 2011). Founder genotyping was through deep sequencing (MiSeq, Ilumina). Mosaic founders were crossed to WT to generate heterozygous F1 offspring, which were also deep sequenced to confirm correctly targeted alleles. *Trem2* H157Y mice were genotyped by qPCR with Custom TaqMan SNP Genotyping assays (Thermo Fisher).

To exclude introduction of unexpected mutation, we performed off-target analysis with two heterozygous F1 mice from each of the two founders (1^#^ and 2^#^) using the online tool CRISPOR (http://crispor.tefor.net/) (Haeussler, Schonig et al., 2016). Three putative sites with top CFD scores above 0.3 were identified and examined by Sanger sequencing (GENEWIZ) of PCR amplification products using extracted genomic DNA.

Our mice were housed in a temperature-controlled environment with a 12-h light–dark cycle and free access to food and water. All animal procedures were approved by the Mayo Clinic Institutional Animal Care and Use Committee (IACUC) and in accordance with the National Institutes of Health Guidelines for the Care and Use of Laboratory Animals.

### Introduction of *Trem2* H157Y mutation to 5xFAD amyloid mouse model

*Trem2* H157Y homozygous mice (Trem2^H157Y/H157Y^) were crossed with 5xFAD mice (The Jackson Laboratory, stock # 34840) to obtain the 5xFAD; Trem2^H157Y/+^ offspring. 5xFAD; Trem2^H157Y/+^ mice were used to setup breeding cages to establish the littermate cohorts with three genotypes including 5xFAD; Trem2^+/+^, 5xFAD; Trem2^H157Y/+^, 5xFAD; Trem2^H157Y/ H157Y^.

### Tissue preparation for immunofluorescence staining or biochemical analyses

Blood samples were collected from mice vena cava after isoflurane induced deep anesthesia and stored at 4°C overnight and subsequently centrifuged at 1000 g for 10 min to collect the supernatant as serum. Mice were transcardinally perfused with 0.01M PBS and the brains were dissected out. Half of the brain was fixed in 4% paraformaldehyde (PFA, Fisher Scientific) for 24 hours followed by dehydration with 30% sucrose (Sigma) for 48 hours. Finally, one hemisphere was embedded in O.C.T. compound (SAKURA) and snap-frozen in liquid nitrogen before cryostat sectioning. The other hemisphere was dissected into cortex, hippocampus, midbrain, and cerebellum which were snap-frozen in liquid nitrogen and stored at −80°C. The cortices were then pulverized and divided into 20-30 mg for RNA extraction and 55-65 mg for protein extraction.

Cortical proteins were extracted sequentially with different lysis buffers. Cortical powder was homogenized in Tris-buffered saline (TBS, Fisher Bioreagents, BP2471-500) supplemented with protease inhibitor (cOmplete, Roche) and phosphatase inhibitor (PhosSTOP, Roche) and subjected to ultracentrifugation at 100,000 g for 1 hour at 4°C. The supernatant was collected as TBS lysate. The pellets were then resuspended in TBSX (TBS plus 1% Triton-X100) supplemented with protease inhibitor and phosphatase inhibitor followed by mild agitation at 4°C for 30 min and centrifuged at 100,000 g at 4°C for 1 hour. Supernatant was collected as TBSX lysate. For amyloid bearing mice, the pellet was further resuspended in 5 M guanidine hydrochloride (GND, Sigma) followed by sonication and centrifuged at 100,000 g for 1 hour at 4°C. The supernatant was collected as GND lysate. Total protein concentration in each lysate was measured (Pierce™ BCA Protein Assay Kit, Cat# 23225) before transferring to 96-well storage plates or 1.5 ml tubes and stored at −80°C until further analysis.

### Immunofluorescence staining, X34 stain and quantification

Embedded hemispheres were coronally sectioned at a 40 µm thickness. Referencing the mouse brain atlas (Paxinos & Franklin, 2013), sections located from AP −1.7 mm to AP −2.06 mm were selected for the following procedures. First, brain slices were blocked in blocking buffer (5% goat serum plus 0.25% Triton in PBS) for 1 hour at room temperature (RT), then incubated overnight in primary antibody solution at 4°C. Slices were then incubated in the Alexa Fluor-conjugated secondary antibodies solution (1:1000, Invitrogen) at RT for 2 hours. The primary antibodies used in this paper include anti-IBA1 (Wako, 019-19741, 1:1000), anti-Aβ (MOAB2, Abcam, ab126649, 1:1000), anti-LAMP1 (Abcam, ab25245, 1:500), and anti-CD68 (Bio-Rad, MCA1957,1:500). Fibrillar Aβ plaque staining used free-floating sections from 5xFAD mouse cohorts. Sections were permeabilized with 0.25% Triton X-100 in PBS and stained with 10 µM X-34 (Sigma, SML1953) in a mixture of 40% ethanol and 0.02M NaOH in PBS as described (Ulrich, Ulland et al., 2018). To assess the plaque associated microglia, IBA1 stain was performed after the X34 stain. To quantify signals of Aβ, X34, IBA1, LAMP1 and CD68, images were taken, stitched using Keyence (BZ-X800) at 20X for the whole slice and analyzed in batch by customized macro coding in Image J with the same setting parameters for all the groups. For X34 and IBA1 co-stain, 30-40 images were taken under Confocal (Zeiss) at 40X with a 0.6 zoom. The number of microglia surrounding plaques within the radius of 30 µm were manually counted. Colocalization of IBA1-and X34 was measured for each plaque in a batch-analysis mode of Image J with customized macro coding. Researchers were blinded to genotypes and groups when performing and quantifying the immunofluorescence staining.

### Primary microglia culture

Cortical cells from pups (p1-p3) were isolated, filtered with 100 um cell strainers (Falcon, 352360), and plated in T75 flasks (Genesee, 25-209) with high-glucose DMEM medium (Gibco, 11965084) containing 10% Fetal Bovine Serum (FBS). Medium was changed to medium containing 25 ng/mL recombinant mouse GM-CSF (Gemini Bio, 300-308P) the next day. Tails from each pup were kept for genotyping. Five days after cell plating, medium in each flask was replaced with fresh GM-CSF-containing medium. On day 9 or 10, microglia were collected by shaking the flasks at 200-220 rpm at RT for ∼20 min, resuspended in non-GM-CSF containing medium, and plated into 6-well plates. After 24 hours, medium from each well was collected as conditioned medium. Cells were lysed with RIPA buffer (Millipore, 20-188) supplemented with protease inhibitor (cOmplete, Roche) and phosphatase inhibitor (PhosSTOP, Roche) followed by mild agitation at 4°C for 30 min and centrifugation at 20,000 g at 4°C for 30 min. Supernatant was collected as RIPA lysate.

### Aβ40, Aβ42, Aβ oligomer, sAPPα, sAPPβ, CTFβ and TREM2 ELIZA

Aβ40 and Aβ42 levels in TBS, TBSX, and GND lysates were determined by ELISA as previously described (Shinohara, Petersen et al., 2013) using an end-specific Aβ monoclonal antibody (13.1.1 for Aβ40 and 2.1.3 for Aβ42) and a HRP-conjugated detection antibody (Ab5, from Dr. Golde lab) (Chakrabarty, Li et al., 2018). Aβ42 in ISF was detected by commercial kits (Thermo Fisher, KHB3544). Aβ oligomers in TBS and TBSX lysates were detected by commercial kits (Biosensis, BEK-2215-2P). sAPPa, sAPPb in TBS lysates were detected by commercial kits (Meso Scale Discovery, K15120E-2). CTFβ in TBSX lysates was detected by commercial kit (IBL, 27776).

TREM2 in TBS lysate, TBSX lysate, serum, conditioned medium, and microglia RIPA lysates were measured as described (Kleinberger et al., 2017) with minor modification using the Meso Scale Discovery (MSD) platform. Streptavidin-coated 96-well plates (MSD, L55SA-2) were blocked overnight at 4°C in blocking buffer (3% bovine serum albumin and 0.05% Tween-20 in PBS). Capture antibody (R&D Systems, BAF1729, 0.25 ug/ml) was applied at RT for 1 hour. Samples were incubated overnight at 4°C with an established dilution in fresh-prepared sample buffer (1% bovine serum albumin and 0.05% Tween-20 in PBS) supplemented with protease inhibitor (cOmplete, Roche). Detection antibody (R&D Systems, MAB1729,) was applied at RT for 1 hour. Sulfo-tag labeled anti rat antibody (MSD, R32AH-5) was applied at RT for 1 hour, and final measurements were made with Read Buffer (MSD, R92TC-3). TBS lysate, TBSX lysate, and serum from Trem2-KO mice were used as negative controls.

### Hippocampal LTP recordings and analyses

Electrophysiological recordings were performed with littermates of *Trem2* H157Y homozygous mice and WT time at 6 months of age as previously described (Rogers, Liu et al., 2017) with minor modifications. Each mouse was acutely decapitated and the brain was dissected out to conduct transverse slicing in ice-cold cutting solution containing 110 mM sucrose, 60 mM NaCl, 3 mM KCl, 1.25 mM NaH2PO4, 28 mM NaHCO3, 0.6 mM sodium ascorbate, 5 mM glucose, 7 mM MgCl2 and 0.5 mM CaCl2. Field excitatory post-synaptic potentials (fEPSPs) were obtained from area CA1 stratum radiatum with the use of a glass microelectrode (2 - 4 mΩ) filled with artificial cerebrospinal fluid (aCSF) containing 125 mM NaCl, 2.5 mM KCl, 1.25 mM NaH2PO4, 25 mM NaHCO3, 25 mM glucose, 1 mM MgCl2 and 2 mM CaCl2. fEPSPs were evoked through stimulation of the Schaffer collaterals using a 0.1 millisecond biphasic pulse delivered every 20 seconds. After a consistent response to a voltage stimulus was established, threshold voltage for evoking fEPSPs was determined and the voltage was increased incrementally every 0.5 - 1 mV until the maximum amplitude of the fEPSP was reached (I/O curve). All other stimulation paradigms were induced at the same voltage, defined as 50-60% of the stimulus voltage used to produce the maximum fEPSP amplitude, for each individual slice. Paired-pulse facilitation (PPF) was induced with two paired-pulses given with an initial delay of 20 milliseconds and the time to the second pulse incrementally increased 20 milliseconds until a final delay of 300 milliseconds was reached. The fEPSP baseline response was then recorded for 20 min. The tetanus used to evoke LTP was a theta-burst stimulation (TBS) protocol consisting of five trains of four pulse bursts at 200 Hz separated by 200 milliseconds, repeated six times with an inter-train interval of 10 seconds. Following TBS, fEPSPs were recorded for 60 min.

All analyses were performed by customized programming in MATLAB (R2019a). The fEPSP slope was calculated within the first 1 ms of the descending domain. I/O curve was presented as the fEPSP slope versus fiber volley amplitude responding to increasing stimulus intensities. PPF strength was examined by the ratio of the second fEPSP slope and first fEPSP slope for each stimulation pair. Potentiation was measured as the increase of the mean fEPSP slope in each minute following TBS normalized to the mean fEPSP descending slope of baseline recordings.

### *In vivo* microdialysis

To assess the Aβ clearance, we examine the Aβ level in hippocampal interstitial fluid (ISF) by *in vivo* microdialysis in awake, free-moving mice as previously described (Cirrito et al., 2003, Liu et al., 2017). Animals were placed in a stereotaxic device equipped with dual manipulator arms and an isoflurane anesthetic mask (David Kopf Instruments). Under isoflurane volatile anesthetic, guide cannula (BR style; Bioanalytical Systems) were cemented into the hippocampus (3.1 mm behind bregma, 2.5 mm lateral to midline, and 1.2 mm below dura at a 12 ° angle). Four to six hours post-surgery, a microdialysis probe (30-kilodalton MWCO membrane, Bioanalytical Systems) was inserted through the guide cannula into the brain. Artificial cerebrospinal fluid (aCSF) (mM: 1.3 CaCl2, 1.2 MgSO4, 3 KCl, 0.4 KH2PO4, 25 NaHCO3, and 122 NaCl, pH 7.4) containing 3% bovine serum albumin (BSA; Sigma) filtered through a 0.1 mm membrane was used as microdialysis perfusion buffer. Flow rate was a constant 1.0 ml/min. Samples were collected hourly into a refrigerated fraction collector. The baseline samples were collected for 10 hours followed by subcutaneous administration of a γ-secretase inhibitor, LY411575 (5 mg/kg) to rapidly block the production of Aβ. Samples were collected for another 4 hours after treatment. ISF Aβ42 in the 14 samples for each mouse was measured by ELISA (Invitrogen, KHB3441, 1:4). To determine Aβ42 half-life (Cirrito et al., 2003), datapoints from drug delivery were analyzed. Meeting with the first-order processes, the elimination rate (*Ke*) of Aβ42 is related to the slope (*a*) of the semi-log plot of concentration versus time: *a* = − *Ke/*2.3. The half-life (T_1/2_) of Aβ42 is further calculated as T1/2 = 0.693/*Ke*.

### Statistical analyses

All data were reported as mean values ± SEM unless. Generally, if sample sizes are larger than 7, to ensure that results were valid in the presence of non-normal distributions, or differing variances between groups, Kruskal-Wallis tests with uncorrected Dun’s multiple comparisons or Wilcoxon Rank-sum tests were used. If the sample size ≤ 7 and dataset showed similar variances examined by F-test, unpaired *t* test was used since nonparametric tests would have very low power. In Fig 4C, unpaired *t* test with Welch’s correction (Welch’s *t* test) was used because of the significant different variances. One-Way ANOCOVA with comparison of slopes was used in Fig 2A. All the statistical analyses were conducted using GraphPad Prism v8.4.3 except for Fig2 in *MATLAB*. All statistical tests were two-sided. The statistical tests used for each analysis, the sample size and the significance levels are reported in the legend of each figure.

## Acknowledgments

This work was supported by NIH grants R01AG066395 (to G.B. and N.Z.), RF1AG056130, R37AG027924, and RF1AG046205 (to G.B).

## Author contributions

WQ, NZ and GB developed the research concept and designed the experiments; WQ, and YC prepared the animals and tissues, and performed most experiments including LTP recording and analysis, immunofluorescence staining/imaging/quantification, Western blotting, ELISA; YAM, WQ, and YC designed and performed primary microglia culture; JAK and C-CL performed the *in vivo* microdialysis; C-CL. coordinated the generation of *Trem2* mouse lines; KA, YC and WQ performed the behavioral tests and analysis; YC maintained the animal colonies and performed genotyping; KC and FL helped with animal tissue collection; FS helped with the image analysis; YAM and MD helped with ELISA; JF supervised the behavioral experiments; WQ, NZ and GB wrote the manuscript with critical inputs and edits by all the co-authors.

## Expanded View Figure legends

**Figure EV1. Analysis of potential off target effects in the *Trem2* H157Y knock-in mice**.

A Top three putative off targets (A) with Cutting Frequency Determination (CFD) Score ranging from 0.25 to 0.44 were identified and sequenced with primers accordingly.

B Single peaks were seen at the putative sites (arrowhead), while two signals were seen at the Trem2 H157Y target site (highlighted with red, arrowhead). Orange arrows indicate the putative region and direction recognized by gRNA.

**Figure EV2. *Trem2* H157Y mutation does not affect microglia density and morphology**.

A Representative images of IBA1 staining are shown for WT, Het, and Hom mice at 6 months of age. Scale, 400 µm.

B-C Cortical microglia (MG) number (B) and cell body size (C) are quantified in Image J for each genotype at 6 months of age. N =11-14 mice per genotype, mixed sex.

D-F Representative confocal images (D) of IBA1 staining were processed (E) and skeletonized (F) in image J for each genotype at 6 months of age. Scale bar for A and B, 50 µm; Scale bar for C 10 µm.

G-I The branch number (G), junction number (H), and total branch length per microglia (MG)

(I) were assessed for each genotype at 6 months of age. N = 9-10 mice per genotype, mixed sex.

Data information: Data are presented as Mean±SEM. Kruskal-Wallis tests with uncorrected Dun’s multiple comparisons were used were used in B-C, G-I. N.S., not significant.

**Figure EV3. *Trem2* H157Y mutation does not affect anxiety, working memory and associative memory**.

A Open field analysis (OFA) was conducted to examine the anxiety of mice with different genotypes at 6 months of age. N =37-40 mice per genotype, mixed sex.

B Y-maze spontaneous alteration test was conducted to examine the working memory of mice with different genotypes at 6 months of age. N =23-26 mice per genotype, mixed sex.

C Contextual fear conditioning test (CFC) was conducted to examine the associative memory of mice with different genotypes at 6 months of age. N =37-40 mice per genotype, mixed sex.

D Cued fear conditioning test (CFC) was conducted to examine the associative memory of mice with different genotypes at 6 months of age. N =37-40 mice per genotype, mixed sex.

Data information: Data are presented as Mean±SEM. Kruskal-Wallis tests with uncorrected Dun’s multiple comparisons were used in A-D. N.S., not significant.

**Figure EV4. Effects of *Trem2* H157Y mutation on Aβ levels, neuronal dystrophy**

A Representative images of fibrillar amyloid staining with X34 are shown for 5xFAD/WT, 5xFAD/Het, and 5xFAD/Hom mice at 8.5 months of age. Scale, 400 µm.

B-C plaque number (B) and size (C) are quantified for each genotype at 8.5 months of age. N =19-24 mice per genotype, mixed sex.

D-E Aβ40 is quantified by ELISA in cortical TBS (A) and TBSX (B) lysates of mice at 8.5 months of age. N = 19-24 mice per genotype, mixed sex.

F-G Aβ42 is quantified by ELISA in cortical TBS (C) and TBSX (D) lysates of mice at 8.5 months of age. N = 19-24 mice per genotype, mixed sex.

H-I Aβ40 is quantified by ELISA in cortical TBS (A) and TBSX (B) lysates of mice from Founder 2# at 8.5 months of age. N = 8-13 mice per genotype, mixed sex.

J-K Aβ42 is quantified by ELISA in cortical TBS (A) and TBSX (B) lysates of mice from Founder 2# at 8.5 months of age. N = 8-13 mice per genotype, mixed sex.

L Representative images of LAMP1 staining are shown for 5xFAD/WT, 5xFAD/Het, and 5xFAD/Hom at the age of 8.5 months of age. Scale, 400 µm.

M LAMP1 immuno-reactivity was quantified for each genotype at the age of 8.5 months of age. N =19-24 mice per genotype, mixed sex.

Data information: Data are presented as Mean±SEM. Kruskal-Wallis tests with uncorrected Dun’s multiple comparisons were used in B-K and M. N.S., not significant. * p<0.05.

**Figure EV5. *Trem2* H157Y mutation does not affect APP processing**.

A-B Soluble APPa (sAPPa, I), Soluble APPβ (sAPPβ, J) were examined, quantified, and normalized to 5xFAD/WT in TBS lysates of mice at 8.5 months of age. N = 19-24 mice per genotype, mixed sex.

C CTFβ was examined, quantified, and normalized to 5xFAD/WT in TBSX lysates of mice at 8.5 months of age. N = 19-24 mice per genotype, mixed sex.

Data information: Data are presented as Mean±SEM. Kruskal-Wallis tests with uncorrected Dun’s multiple comparisons were used in A-C. N.S., not significant.

